# Maintenance of epithelial traits and resistance to mesenchymal reprogramming promote proliferation in metastatic breast cancer

**DOI:** 10.1101/2020.03.19.998823

**Authors:** Laura Eichelberger, Massimo Saini, Helena Domínguez Moreno, Corinna Klein, Johanna M. Bartsch, Mattia Falcone, Manuel Reitberger, Elisa Espinet, Vanessa Vogel, Elisabeth Graf, Thomas Schwarzmayr, Tim-Matthias Strom, Mareike Lehmann, Melanie Königshoff, Nicole Pfarr, Roberto Würth, Elisa Donato, Simon Haas, Saskia Spaich, Marc Sütterlin, Andreas Schneeweiss, Wilko Weichert, Gunnar Schotta, Andreas Trumpp, Martin R. Sprick, Christina H. Scheel

**Affiliations:** Institute of Stem Cell Research, Helmholtz Center Munich, Neuherberg, Germany; Heidelberg Institute for Stem Cell Technology and Experimental Medicine (HI-STEM gGmbH), Heidelberg, Germany; Division of Stem Cells and Cancer, German Cancer Research Center (DKFZ), and DKFZ-ZMBH Alliance, Heidelberg, Germany; Department of Biomedicine, University Basel and University Hospital Basel, Basel, Switzerland; Biomedical Center (BMC) Munich, Ludwig-Maximilians-University (LMU) Munich, Planegg-Martinsried, Germany; Institute of Human Genetics, Helmholtz Center Munich, Neuherberg, Germany; Research Unit Lung Repair and Regeneration, Helmholtz Center Munich, Member of the German Center of Lung Research (DZL), Munich, Germany; Institute of Pathology, Technical University Munich (TUM), Munich, Germany; Department of Gynaecology and Obstetrics, University Women’s Clinic, University Medical Centre Mannheim, Mannheim, Germany; National Center for Tumor Diseases, University Hospital and German Cancer Research Center, Heidelberg, Germany; Department of Pathology, Heidelberg University Hospital, Heidelberg, Germany; German Cancer Consortium (DKTK); Department of Dermatology, St. Josef Hospital, Ruhr-University Bochum, Bochum, Germany

**Keywords:** EPCAM, ZEB1, EMT, resistance, breast cancer, metastasis

## Abstract

Despite important advances in the treatment of breast cancer, the 5-year survival rate for patients with distant metastasis remains less than 30%. Metastasis is a complex, multi-step process beginning with local invasion and ending with the outgrowth of systemically disseminated cells into actively proliferating metastases that ultimately cause the destruction of vital organs. It is this last step that limits patient survival and, at the same time, remains the least understood mechanistically. Here, we focus on understanding determinants of metastatic outgrowth using metastatic effusion biopsies from stage IV breast cancer patients. By modelling metastatic outgrowth through xenograft transplantation, we show that tumour initiation potential of patient-derived metastatic breast cancer cells across breast cancer subtypes is strongly linked to high levels of EPCAM expression. Breast cancer cells with high EPCAM levels are highly plastic and, upon induction of epithelial-mesenchymal transition (EMT), readily adopt mesenchymal traits while maintaining epithelial identity. In contrast, low EPCAM levels are caused by the irreversible reprogramming to a mesenchymal state with concomitant suppression of metastatic outgrowth. The ability of breast cancer cells to retain epithelial traits is tied to a global epigenetic program that limits the actions of EMT-transcription factor ZEB1, a suppressor of epithelial genes. Our results provide direct evidence that maintenance of epithelial identity is required for metastatic outgrowth while concomitant expression of mesenchymal markers enables plasticity. In contrast, loss of epithelial traits is characteristic of an irreversible mesenchymal reprogramming associated to a deficiency for metastatic outgrowth. Collectively, our data provide a framework for the intricate intercalation of mesenchymal and epithelial traits in metastatic growth.

## Introduction

The outgrowth of macrometastases in vital organs causes almost all breast cancer deaths^1^. Previously, it was shown that circulating tumour cells (CTCs) in blood of breast cancer patients expressing the Epithelial Cell Adhesion Molecule (EPCAM) are capable of initiating metastatic tumours *in vivo*^2^. However, it has been questioned whether EPCAM is the most useful marker for CTCs since its expression is downregulated when breast cancer cells undergo an epithelial-mesenchymal transition (EMT)^3^.

Experimental studies addressing the impact of EMT in different cancers have yielded a wealth of controversial results. For example, it has been shown that the expression of the EMT-inducer *Twist1* decreases proliferation of tumour cells, but increases their abundance within the circulation^4^. Moreover, recent findings in a mouse model of pancreatic carcinoma show that deletion of *Zeb1*, another EMT-transcription factor, decreases invasion and lung colonization without affecting primary tumour growth, suggesting that Zeb1 is crucial for tumour progression^5^.

Other studies emphasize that a transient, rather than permanent expression of EMT-transcription factors is crucial for the outgrowth of metastases^4,6,7^. This is supported by the finding that most macroscopic metastases generated by carcinomas display an epithelial morphology^8^. However, a strict requirement for EMT at any time during the metastatic process is called into question by the finding that overexpression of micro RNAs that inhibit EMT does not affect metastasis^9^ and loss of *Twist1* and *Snai1* expression in a pancreatic cancer mouse model does not change invasion and disease progression^10^.

More complication has recently arisen from the observation that tumour cells with an intermediate, often termed hybrid epithelial-mesenchymal phenotype, are the most competent in colonization and metastasis formation^11^.

Some of this apparent controversy might be due to differences in the tissue- or cell-of cancer origin. Nonetheless, the fundamental question how different phenotypes, i.e. epithelial, mesenchymal, or hybrids arise, and how they contribute to metastatic outgrowth remains poorly understood. To address this question in the context of breast cancer, we examined the *in vivo* tumour-propagating ability of cancer cells derived from effusion biopsies of metastatic breast cancer patients. We observed that, across different breast cancer subtypes, EPCAM^high^ cells were consistently more efficient in establishing tumours in immunocompromised mice compared to EPCAM^low^ cells. These results guided subsequent mechanistic studies to understand why EPCAM^low^ cells from metastatic biopsies failed to propagate the disease. Thereby we discovered that a global epigenetic program prevents repression of epithelial genes in EPCAM^high^ cells upon challenge by an EMT-inducing stimulus. Thus, our data suggest that the ability to display epithelial-mesenchymal plasticity while maintaining epithelial identity promotes metastatic outgrowth.

## Results

### High EPCAM levels in patient-derived metastatic breast cancer cells correlate with tumour propagation capacity

To determine whether distinct tumour cell phenotypes contribute differently to metastatic outgrowth, we used xenograft transplantation of cells from human cancers as a proxy assay. For this purpose, we analysed liquid biopsies (pleural and ascitic effusions) from stage IV breast cancer patients encompassing all major subtypes of breast cancer (Supplementary Figure 1a). After excluding immune cells using CD45, we identified three subpopulations of tumour cells present in these biopsies according to cell surface levels of the Epithelial Cell Adhesion Molecule (EPCAM): EPCAM^high^, EPCAM^low^, and EPCAM^neg^ (Figure 1a, Supplementary Figure 1b). Next, we assessed the tumour-initiating capacity (TIC) of each subpopulation *in vivo* by orthotopic implantation in immunocompromised NSG mice to generate patient-derived xenografts (PDX, Figure 1b). Strikingly, limiting dilution analysis revealed that across all subtypes, EPCAM^high^ cells were 30 times more efficient in generating tumours (TIC frequency: 6.3×10^−4^) compared to EPCAM^low^ cells (2.1×10^−5^). EPCAM^neg^ cells showed the lowest efficiency in tumour generation (1×10^−6^, Figure 1b). Importantly, the PDX-outgrowths recapitulated the subtype of the primarily diagnosed tumour, as shown by subtype-specific marker expression using immunohistochemistry (Supplementary Figure 1c). Additionally, metastatic nodules in the lungs were exclusively found in mice injected with EPCAM^high^ cells (Figure 1b) and again expressed markers congruent with the primary cancer subtype (Supplementary Figure 1d). Thus, these data suggest that EPCAM^high^ patient-derived breast cancer cells from metastatic sites preferentially contribute to outgrowth compared to EPCAM^low^ and EPCAM^neg^ cells.

**Figure 1:**
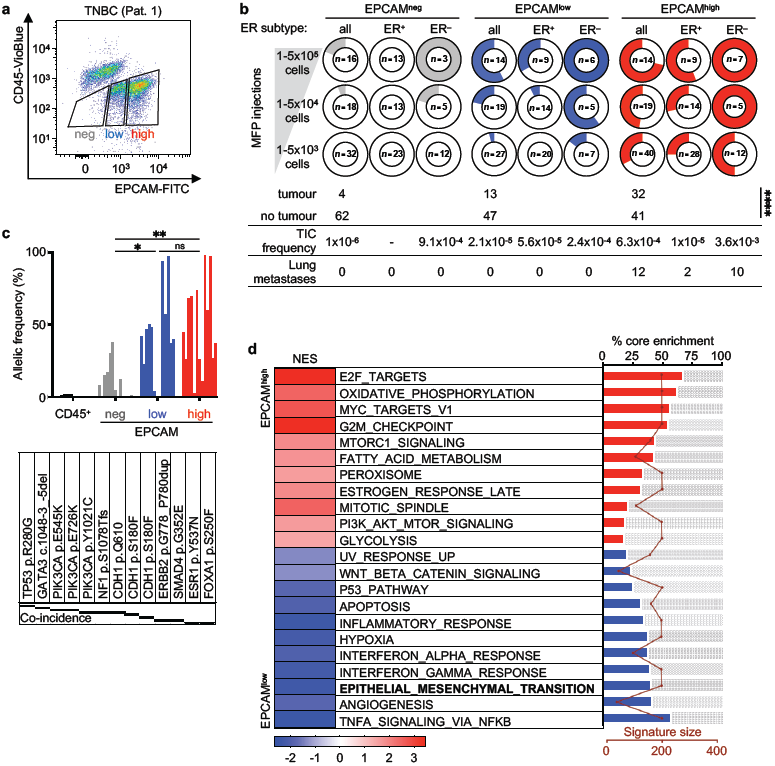
High EPCAM expression correlates with disease-propagating potential in metastatic breast cancer cells. **a**, FACS plot of CD45 and EPCAM of cells from liquid biopsies. Gates used for sorting cells with different EPCAM levels are highlighted. TNBC, Triple Negative Breast Cancer; Pat., Patient. **b**, Number of mice with tumours formed by cells injected into the mammary fat pad in limiting dilution. The number of mice injected (n=x) is indicated at the centre of each chart. The table shows numbers of mice with orthotopic tumour growth (tumour) and no orthotopic tumour growth (no tumour), the calculated TIC frequency of tumour-initiating cells (TIC), and the number of mice with lung metastases. Chi-square test; p-value: ****<0.0001. ER, oestrogen receptor. **c**, Allelic frequency values in sorted CD45^+^, EPCAM^neg^ (neg), EPCAM^low^ (low), and EPCAM^high^ (high) cell subsets presented as a single percentage value per mutation per patient. Mutations are ordered as bars from left to right as they are listed in the table below. n=7; Wilcoxon matched-pairs signed rank test; p-values: *=0.039, **=0.0098, ns=not significant. **d**, Hallmark gene set enrichment analysis of RNA-sequencing data of sorted EPCAM^high^ and EPCAM^low^ cells from an ER^+^ and an ER^−^ patient. Signatures with FDR<0.25 are ranked by the % of core-enrichment. Signature size indicates the overall number of genes composing the reference signature. n=4.

To take a closer look at breast cancer subtype-specific differences, effusion biopsies were stratified according to oestrogen receptor (ER) status. Cells from patients with ER-positive tumours (ER^+^) showed the highest TIC frequency within the EPCAM^high^ population (1×10^−5^) compared to EPCAM^low^ (5.6×10^−5^), while the EPCAM^neg^ population lacked detectable TICs (Figure 1b). In biopsies from patients with ER-negative tumours (ER^−^), all populations initiated tumours to a certain extent (Figure 1b), consistently with a higher degree of phenotypic plasticity previously observed in ER^−^ liquid biopsies^12^. Nonetheless, we observed a 15 times higher TIC frequency in EPCAM^high^ cells (3.6×10^−3^) compared to EPCAM^low^ cells (2.4×10^−4^), and the lowest TIC frequency in EPCAM^neg^ cells (9.1×10^−4^, Figure 1b).

Since metastatic effusion biopsies are derived from body cavities lined with mesothelium, contamination with sloughed-off, normal mesothelial cells could potentially provide a simple explanation for the lower TIC frequency in EPCAM^low^ and EPCAM^neg^ cell populations. To test this possibility, we assessed the mutational spectrum in EPCAM-sorted subpopulations by using panel-based targeted sequencing covering 50 commonly mutated breast cancer genes. Thus, 13 different somatic mutations were detected and their allelic frequency evaluated in each subpopulation (Figure 1c). This analysis revealed that across subtypes, the frequency of mutated alleles in the EPCAM^neg^ population was significantly lower compared to the EPCAM^low^ (p=0.039) or EPCAM^high^ populations (p=0.0098, Figure 1c), indicating that the EPCAM^neg^ population contained a significant amount of non-tumour cells. By contrast, EPCAM^high^ and EPCAM^low^ populations showed comparable allelic frequencies for all mutations detected.

Together, these results suggest that the observed differences in TIC frequency in the EPCAM^neg^ population might arise from contamination with normal cells, whereas the observed differences between EPCAM^high^ and EPCAM^low^ populations originate from qualitative differences between these populations. Since we observed highly similar mutation profiles between these populations, we speculated that the observed differences in TIC frequency might arise from differences in epigenetic rather than genetic cellular programs.

To further explore this hypothesis, we performed RNA-sequencing of freshly sorted EPCAM subpopulations from an ER^+^ and an ER^−^ patient. Among larger and non-redundant gene set collections, one of the most enriched features in EPCAM^low^ cells was the EMT process (Figure 1d, Supplementary Figure 1e). By contrast, EPCAM^high^ cells were enriched for multiple features associated with an active cell cycle, oxidative phosphorylation, MYC targets and in general, pathways associated with cell growth (Figure 1d, Supplementary Figure 1f).

This result supported the conclusion derived from xenograft transplantation: in metastatic effusion biopsies, EPCAM expression serves as a marker distinguishing between disease-propagating, actively growing tumour cells and less proliferative subpopulations. Moreover, RNA-sequencing pointed us towards EMT programs as a potential underlying mechanism for the reduction in TICs we observed in the EPCAM^low^ cells.

### Contrasting responses to an EMT-inducing stimulus generate morphological and functional heterogeneity in breast cancer cells

To investigate whether EMT programs in breast cancer cells contribute to a suppression of metastatic outgrowth and to explore the role of epithelial traits, we induced EMT in epithelial breast cancer cells under defined conditions. To do so, we used immortalized human mammary epithelial cells (HMLE) engineered to express a conditional *Twist1*-ER (HMLE-Twist1-ER) activatable by Tamoxifen (TAM) treatment^7,13^.

After 21 days of persistent Twist1-activation all cells, based on morphology, appeared to have trans-differentiated from an epithelial to a mesenchymal phenotype (Supplementary Figure 2a). Indeed, analysis of cell-surface EPCAM expression after Twist1-activation revealed that most cells had lost EPCAM cell surface expression. However, a minor subset of cells continued to express EPCAM (Figure 2a, Supplementary Figure 2c), indicating that the latter subset of cells was resistant to EMT-induction. To functionally address this heterogeneity, we prospectively sorted EPCAM^neg^ and EPCAM^pos^ cells (Figure 2a). Sorted cells were transferred to Tamoxifen-free conditions for 10 days (+TAM 21d/-TAM 10d) to reproduce the effect of a transient EMT stimulus, as it is thought to take place during tumour progression^14^. Based on morphology and expression of markers such as E-cadherin (*CDH1*), EPCAM^neg^ cells could be identified as a population that remained mesenchymal even after cessation of Twist1-activation. By contrast, EPCAM^pos^ cells displayed an epithelial phenotype, thus showing insensitivity to the EMT-inducing stimulus provided by Twist1-activation (Figure 2b, c, Supplementary Figure 2b). Next, we tested the ability of EPCAM^pos^ and EPCAM^neg^ cells to generate organoids in a three-dimensional collagen gel-based organoid assay to mimic more physiological growth conditions^7,15^. Strikingly, only EPCAM^pos^ cells proliferated, whereas EPCAM^neg^ cells failed to generate organoids (Figure 2d, e).

**Figure 2:**
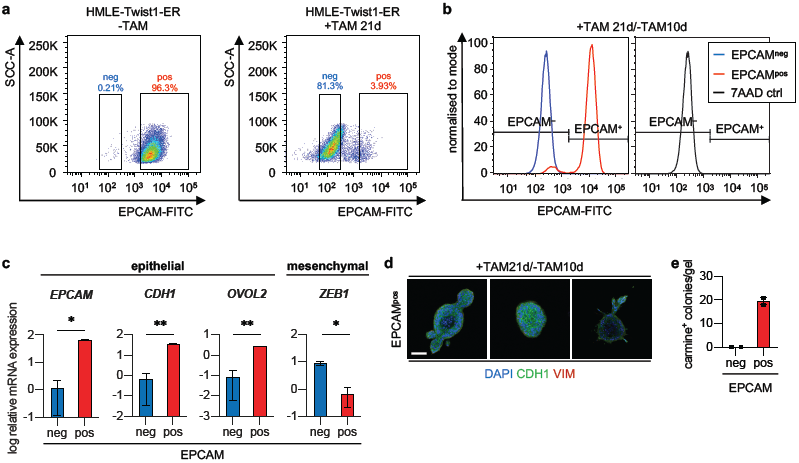
A subset of HMLE-Twist1-ER cells maintains EPCAM expression during EMT-induction. **a**, FACS plot of EPCAM of HMLE-Twist1-ER bulk cells untreated (-TAM) or treated with Tamoxifen for 21 days (+TAM 21d). The gates used for cell sorting are highlighted. **b**, Flow cytometric staining of EPCAM of sorted EPCAM^neg^ and EPCAM^pos^ cells after 10 days of Tamoxifen withdrawal (+TAM21d/-TAM10d). Gates for EPCAM were set according to unstained control as indicated by cells only stained for 7AAD (7AAD ctrl). **c**, log relative mRNA expression levels of *EPCAM, CDH1, OVOL2*, and *ZEB1* of sorted EPCAM^neg^ (neg) and EPCAM^pos^ (pos) HMLE-Twist1-ER cells treated as described in b. n=2; mean ± SEM; multiple t-tests (Holm-Sidak correction); p-values: *<0.05, **<0.005. **d**, Immunofluorescence staining of DAPI (blue), E-cadherin (CDH1, green), and Vimentin (VIM, red) of organoids formed by sorted EPCAM^pos^ HMLE-Twist1-ER cells in floating 3D-collagen gels treated as described in b. Scale bar: 50 µm. Positive control for Vimentin not shown. **e**, Number of carmine positive-stained colonies per gel of sorted EPCAM^neg^ (neg) and EPCAM^pos^ (pos) HMLE-Twist1-ER cells in floating 3D-collagen gels treated as described in b. n=2; mean ± SEM.

Together, these results showed that contrasting cellular responses to an EMT-inducing stimulus create heterogeneity with respect to EPCAM-expression in a breast cancer cell line model similar to what we encountered in metastatic effusion biopsies (Figure 1). Moreover, these data suggest that a proportion of breast cancer cells can resist induction of EMT and maintain epithelial properties, whereas cells that undergo irreversible EMT lose their ability to proliferate in a three-dimensional environment.

### Single-cell level analysis confirms that breast cancer cells either intrinsically resist EMT-induction or enter a stable, mesenchymal cell state

Based on these data, we set out to understand why some cells appeared to be more susceptible to EMT than others. For this purpose, we generated several single-cell clones (SCCs, Supplementary Figure 3a) from the HMLE-Twist1-ER bulk population and treated the clones with TAM independently to assess the competence for EMT at the single-cell level. Again, we observed that only a subset of single-cell clones (SCCs) underwent EMT (42 of 58, 72%), indicated by a gain of mesenchymal traits and the concurrent loss of epithelial traits (M-SCCs: M1-3, Figure 3a, b). The remaining SCCs (16 of 58, 28%) failed to undergo EMT and remained epithelial (E-SCCs: E1-3, Figure 3a, b). Importantly, Twist1 nuclear localisation and expression levels, as well as transcript levels of the direct target gene *WNT5A*^16^ were quantitatively comparable in all clones (Supplementary Figure 3b-d), ruling out variable Twist1 expression levels as the reason for the differences in susceptibility for EMT induction.

**Figure 3:**
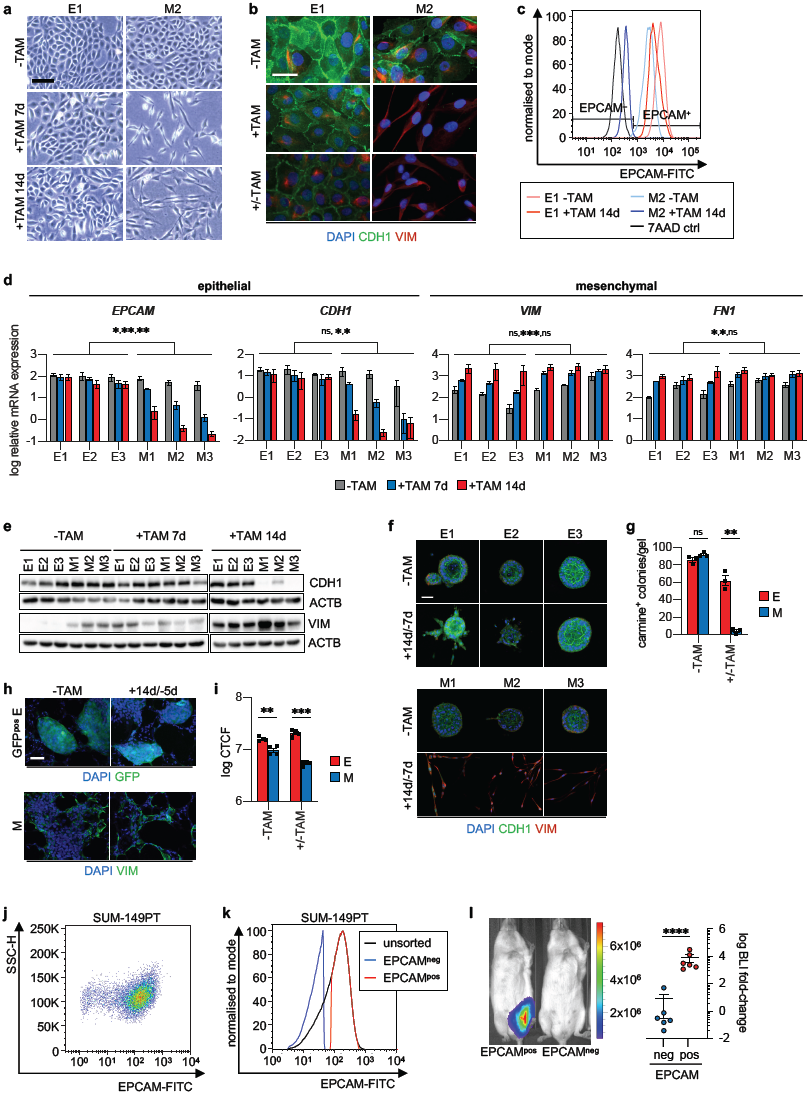
A subset of breast cancer single-cell clones resists complete EMT and maintains the ability to proliferate in different environments. **a**, Bright field images of a representative E-SCC (E1) and a representative M-SCC (M2), untreated (-TAM), treated with Tamoxifen for 7 days (+TAM 7d), or treated with Tamoxifen for 14 days (+TAM 14d). Scale bar: 100 µm. **b**, Immunofluorescence staining of DAPI (blue), E-cadherin (CDH1, green), and Vimentin (VIM, red) of a representative E-SCC (E1) and a representative M-SCC (M2), untreated (-TAM), treated with Tamoxifen for 15 days (+TAM), or treated with Tamoxifen for 15 days followed by Tamoxifen withdrawal for 9 days (+/-TAM). Scale bar: 20 µm. **c**, Flow cytometric staining of EPCAM of a representative E-SCC (E1) and a representative M-SCC (M2), untreated (-TAM) or treated with Tamoxifen for 14 days (+TAM 14d). Gates for EPCAM were set according to a control as indicated by control cells only stained for 7AAD (7AAD ctrl). **d**, log relative mRNA expression levels of *EPCAM, CDH1, VIM*, and *FN1* in E-SCCs (E1-E3) and M-SCCs (M1-M3), treated as described in a. n=3; mean ± SEM; multiple t-tests (Holm-Sidak correction); p-values: *<0.05, **<0.005, ***<0.0005, ns=not significant. **e**, Immunoblot of E-cadherin (CDH1), Vimentin (VIM), and β-actin (ACTB) of E-SCCs (E1-E3) and M-SCCs (M1-M3), treated as described in a. **f**, Immunofluorescence staining of DAPI (blue), E-cadherin (CDH1, green), and Vimentin (VIM, red) of E-SCCs (E1-E3) and M-SCCs (M1-M3), untreated (-TAM) or after 14 days of Tamoxifen treatment followed by Tamoxifen withdrawal for 7 days (+14d/-7d) growing in floating 3D-collagen gels. **g**, Number of carmine positive stained colonies per gel of E-SCCs and M-SCCs treated as described in f. n=3; mean ± SEM; multiple t-tests (Holm-Sidak correction); p-values: **=0.002, ns=not significant. **h**, Immunofluorescence staining of DAPI (blue) of GFP^pos^ E-SCC (E) cells (green) and of DAPI (blue) and Vimentin (VIM, green) of M-SCC (M) cells growing on PCLS, untreated (-TAM) or after 14 days of Tamoxifen treatment followed by 5 days of Tamoxifen withdrawal (+/-TAM). Scale bar: 50 µm. **i**, log CTCF of DAPI intensity of representative images of GFP^pos^ E-SCC (E) cells and M-SCC (M) cells growing on PCLS treated as described in h. n=4; mean ± SEM; multiple t-tests (Holm-Sidak correction); p-values: **=0.0022, ***=0.0001. CTCF, Corrected Total Cell Fluorescence. **j**, Flow cytometric staining of EPCAM of SUM-149PT bulk cells. **k**, Flow cytometric staining of EPCAM of unsorted bulk, sorted EPCAM^neg^, and EPCAM^pos^ SUM-149PT cells. **l**, Bioluminescence imaging of NSG mice after intra-femoral injection of an equal number (10,000) of EPCAM^neg^ (neg) and EPCAM^pos^ (pos) luciferase-labelled SUM-149PT cells showing fold-change to baseline signal at the termination of the experiment (20 weeks post-injection). The image shows two representative mice measured 10 weeks post-injection. The colour scale indicates the radiance of photons/sec/cm^2^/sr. n=6; mean ± SEM; unpaired student’s t-test with Welch’s correction; p-value: ****<0.0001. BLI, bioluminescence intensity.

In detail, Twist1-activation led to a complete loss of surface EPCAM expression in M-SCCs, while we observed only a minor shift in E-SCCs (Figure 3c, d). Moreover, expression of E-cadherin (CDH1) was maintained in E-SCCs, but not M-SCCs (Figure 3b, d, e). By contrast, expression of mesenchymal markers such as Vimentin (VIM) was upregulated upon Twist1-activation in both E-SCCs and M-SCCs to a similar extent (Figure 3b, d, e). Thus, E-SCCs adopted a transient hybrid phenotype during Twist1-activation, characterized by concomitant expression of epithelial and mesenchymal markers.

By contrast, epithelial markers were permanently downregulated in M-SCCs with concomitant high expression levels of mesenchymal genes (Supplementary Figure 3e, f). Thus, M-SCCs did not undergo a mesenchymal-epithelial transition (MET) following inactivation of Twist1, but persisted in a mesenchymal state.

Importantly, the contrasting responses to Twist1 in E-SCCs and M-SCCs did not simply reflect differences in kinetics to EMT-induction: during prolonged Twist1-activation for 21 or 28 days, E-SCCs still maintained an epithelial phenotype and the expression of epithelial markers such as E-cadherin (*CDH1)* (Supplementary Figure 3g, h). This result suggests a strong protection of epithelial traits in E-SCCs that cannot be overcome by long-term activation of an EMT-inducing stimulus.

### Resistance to complete EMT correlates with the ability of breast cancer cells to proliferate in a three-dimensional environment

Next, we wished to determine whether resistance to EMT again correlated with the capacity for growth in a three-dimensional environment. To do so, individual SCCs were plated in a collagen gel-based organoid assay^7,15^. Thereby, we observed that prior to an EMT-stimulus E-SCCs and M-SCCs formed epithelial organoids expressing E-cadherin with similar frequency (Figure 3f, g). However, when E-SCCs and M-SCCs were plated into the organoid assay after 14 days of transient Twist1-activation, only E-SCCs generated epithelial organoids, as observed in breast cancer metastases^8^. By contrast, M-SCCs remained in the collagen gel as single cells and failed to initiate proliferation (Figure 3f, g).

To further explore this deficiency in proliferation and approximate conditions during metastatic outgrowth, we tested the growth of E-SCCs and M-SCCs on precision-cut lung slices (PCLS), which allow analysis of 3D-lung tissue architecture *ex vivo*^17^. For this purpose, PCLS were incubated for 5 days in Tamoxifen-free conditions with GFP^pos^ E-SCCs or M-SCCs that were either untreated or treated with TAM for 14 days. Strikingly, even before Twist1-activation, E-SCCs were 1.6-fold more efficient in colonizing the PCLS (Figure 3h, i). However, after transient Twist1-activation, the ability of M-SCCs to proliferate on the lung slices decreased further and E-SCCs colonized the PCLS 3.9-fold more efficient than M-SCCs (Figure 3h, i).

Together, these data further supported our observation that resistance to complete EMT through maintenance of epithelial features promotes the ability of breast cancer cells to initiate proliferation in different three-dimensional environments, possibly correlating with the capacity for metastatic outgrowth.

Since HMLE cells are not tumorigenic in immunocompromised mice we prospectively sorted the EPCAM^pos^ and EPCAM^neg^ subpopulations of SUM-149PT breast cancer cells (Figure 3j, k) to evaluate their *in vivo* metastatic potential. For this, we injected these populations into the hindleg femurs of NSG mice as previously described^2^ in order to evaluate metastasis-initiating activity in the bone, the most frequent site of metastasis in breast cancer. Indeed, EPCAM^pos^ cells generated bone metastases after 20 weeks of observation, while EPCAM^neg^ cells failed to produce metastases (Figure 3l).

Together, data from different experimental models *in vitro* and *in vivo* support the idea that the occurrence of complete EMT negatively impacts metastatic outgrowth whereas the maintenance of epithelial features promotes metastasis.

### Global chromatin changes during transient EMT-induction define EMT-resistance versus-susceptibility

To investigate the mechanistic basis for EMT-resistance versus-susceptibility, we again turned to E-SCCs and M-SCCs derived from HMLE-Twist1-ER cells. Specifically, we wished to determine whether the contrasting responses of SCCs to Twist1-activation could be observed at an epigenetic regulatory level. For this purpose, we performed ATAC-sequencing in conjunction with RNA-sequencing in three E-SCCs and three M-SCCs before Twist1-activation (1), after 7 (2) and 14 days of Twist1-activation (3), and after 14 days of Twist1-activation followed by 7 days of Twist1-deactivation (4).

Principal component analysis (PCA) revealed a comparable landscape of chromatin accessibility in E-SCCs and M-SCCs before Twist1-activation (Figure 4a). However, both at 7 and 14 days after Twist1-activation, M-SCCs harboured the greatest variation in chromatin accessibility compared to all other conditions, which was evident in changes that created a shift on the PC1 axis (Figure 4a). In contrast, during Twist1-activation, changes observed in E-SCCs created a shift on the PC2 axis, corresponding to comparatively minor variance (Figure 4a).

**Figure 4:**
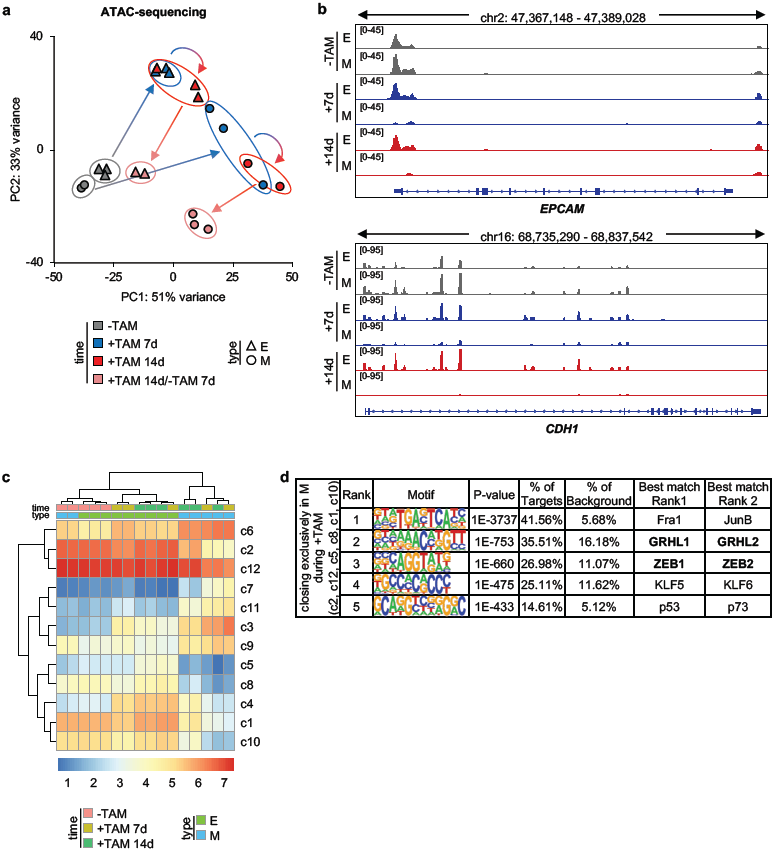
EMT-induction causes genome-wide chromatin and transcriptional changes. **a**, Principal component (PC) analysis of ATAC-sequencing data of E-SCCs (△) and M-SCCs (**◯**), untreated (-TAM), treated with Tamoxifen for 7 days (+TAM 7d), treated with Tamoxifen for 14 days (+TAM 14d), or treated with Tamoxifen for 14 days followed by Tamoxifen withdrawal for 7 days (+TAM 14d/-TAM 7d). Each data point represents one SCC at the indicated time point. **b**, Genome browser high resolution screenshot of ATAC-sequencing data of *EPCAM* and *CDH1* of one representative E-SCC (E) and one representative M-SCC (M), untreated (-TAM), treated with Tamoxifen for 7 days (+7d), or treated with Tamoxifen for 14 days (+14d). **c**, Heatmap of chromatin accessibility of 12 clusters of ATAC-sequencing peaks of E-SCCs and M-SCCs treated as described in b. **d**, Top 5 hits of Homer *de novo* transcription factor motif analysis of grouped clusters closing exclusively in M-SCCs (M) during Tamoxifen treatment (+TAM).

Remarkably, upon Twist1 deactivation (+TAM 14d/-TAM 7d), changes in chromatin accessibility were reversed in E-SCCs but not in M-SCCs (Figure 4a). Thus, the dynamics of chromatin accessibility in E-SCCs and M-SCCs mirrored our observations that during transient Twist1-activation, E-SCCs can temporarily acquire a hybrid phenotype, whereas M-SCCs remain in a stable mesenchymal state. This was further corroborated by focusing on the loci of epithelial genes such as *EPCAM* and *CDH1*: here, we observed that before Twist1-activation, both loci were similarly accessible. However, during Twist1-activation, access to both loci was lost in M-SCCs, but not E-SCCs (Figure 4b). RNA-sequencing revealed comparable dynamics with respect to changes in transcript abundance (Supplementary Figure 4a), indicating that changes observed at the transcript level can be traced back to chromatin accessibility and *vice versa*.

Together, these results reveal that global changes in chromatin accessibility developed during Twist1-activation are less dramatic and reversible in E-SCCs compared to M-SCCs. These changes in chromatin accessibility correlate with differences in gene expression and, as a corollary, the functional differences we observed between E-SCCs and M-SCCs after transient Twist1-activation.

In order to pinpoint transcription factors (TFs) involved in these transitions, we performed peak clustering of the ATAC-sequencing data. By doing so, we confirmed close clustering of E-SCCs and M-SCCs before Twist1-activation. After Twist1-activation, E-SCCs continued to cluster closer with the untreated condition, whereas treated M-SCCs formed a distinct cluster (Figure 4c). Detailed analysis revealed three different types of changes in chromatin accessibility during Twist1-activation: loci previously not accessible that become accessible in both M-SCCs and E-SCCs (c3, c4), loci becoming accessible only in M-SCCs, but not E-SCCs (c6, c7, c11, c9), and loci losing accessibility in M-SCCs, but not E-SCCs (c2, c12, c5, c8, c1, c10, Figure 4c). Of note, the latter clusters contained the loci of epithelial genes, such as *CDH1* and *EPCAM*. Homer *de novo* TF motif analysis of combined clusters revealed specific motifs representing each cluster type (Figure 4d, Supplementary Figure 4b).

Importantly, the results included several motifs that have been previously shown to be enriched in cells undergoing EMT in mice *in vivo*^11^. For example, this included loss of accessibility in M-SCCs for a p53/p73/p63 binding motif, a gain in accessibility for basic leucine zipper TF motifs in M-SCCs, and TEAD and AP-1 motifs as well as a TCF4 binding motif becoming newly accessible upon Twist1-activation in both M-SCCs and E-SCCs (Figure 4d, Supplementary Figure 4b). Interestingly, TCF4 binds to an E-box motif and was shown to interact with TWIST1^18,19^. Additionally, the same cluster type revealed enrichment of a consensus Twist binding motif (Supplementary Figure 4b). The observation that Twist motifs enrich in loci commonly accessible in both E-SCCs and M-SCCs is in line with our observations that Twist1 appeared to be similarly active in E-SCCs and M-SCCs with respect to specific target loci that are actively transcribed (e.g. *WNT5A*).

Together, these data suggest that Twist1 is indeed active in both E-SCCs and M-SCCs and that the different cellular responses to Twist1 activation are not merely the result of Twist1 not functioning as a transcription factor in these cells. As a corollary, our results indicate that the events leading to EMT-resistance or -susceptibility may occur downstream of Twist1-binding to chromatin.

To further investigate this hypothesis, we focused on regions where chromatin accessibility was lost in M-SCCs but not E-SCCs during Twist1-activation (c2, c12, c5, c8, c1, c10, Figure 4c). Here, *de novo* binding motif analysis revealed an enrichment for ZEB1 and ZEB2 motifs (Figure 4d). ZEB1 is known as a key EMT-transcription factor downstream of Twist1 that represses the expression of epithelial genes, such as E-cadherin (*CDH1*)^20^ and *EPCAM*^21^. Additionally, we observed enrichment of GRHL1 and GRHL2 motifs in clusters that become accessible exclusively in M-SCCs during Twist1-activation (Figure 4d). Importantly, grainy-head like factors (GRHL1-3) are known to prime epithelial enhancers for transcriptional activation^22^. Moreover, *GRHL2* expression is controlled by ZEB1 in a negative feedback-loop^23^.

Together, these results suggest ZEB and/or GRHL as mediators determining whether breast cancer cells undergo irreversible EMT or resist trans-differentiation.

### ZEB1 is necessary, but not sufficient to overcome EMT-resistance in breast cancer cells

Next, we looked more closely at the RNA-sequencing results that were obtained concomitantly with ATAC-sequencing data at different time points of Twist1-activation. Expression of *GRHL1* and *GRHL2* was downregulated during EMT in M-SCCs but not E-SCCs. Specifically, high expression levels of *GRHL2* were maintained in E-SCCs, but expression was strongly downregulated in M-SCCs (Figure 5a). In support of our data, GRHL2 was recently shown to be a pioneering transcription factor that maintains the chromatin of epithelial genes in an open state^22,24^. Moreover, when analysing expression levels of *ZEB1* and *ZEB2*, we also observed a clear difference between E-SCCs and M-SCCs that was particularly pronounced for *ZEB1* (Figure 5b, d). With respect to kinetics, *ZEB1* was upregulated in E-SCCs after 14 days of Twist1-activation, whereas in M-SCCs an upregulation could already be observed after 7 days (Figure 5b, d). Additionally, after 14 days of Twist1-activation, *ZEB1* expression levels were 2.9-fold higher in M-SCCs compared to E-SCCs (Figure 5c). These data were in line with the observation that *ZEB1* expression was also higher in sorted EPCAM^neg^ HMLE-Twist1-ER cells compared to sorted EPCAM^pos^ cells (Figure 2c). Next, we determined the expression of ZEB1 downstream targets such as *OVOL2* and *MIR200*-family members, which function in negative feedback loops with ZEB1^20,25,26^. Indeed, transcription of both *OVOL2* and *MIR200*-family members was downregulated in M-SCCs, but not E-SCCs during transient Twist1-activation, suggesting that negative feedback loops restricting ZEB1-activity remained active in E-SCCs, but were disabled in M-SCCs (Figure 5c, Supplementary Figure 5a). In line with these data, epithelial ZEB1-target genes such as *OVOL2* and *CDH1* remained repressed after transient Twist1-activation in M-SCCs, whereas the expression of mesenchymal genes such as *VIM* and *FN1* remained high (Supplementary Figure 3f). Strikingly, prolonged Twist1-activation could not break these feedback loops and overcome EMT-resistance in E-SCCs (Supplementary Figure 3h).

**Figure 5:**
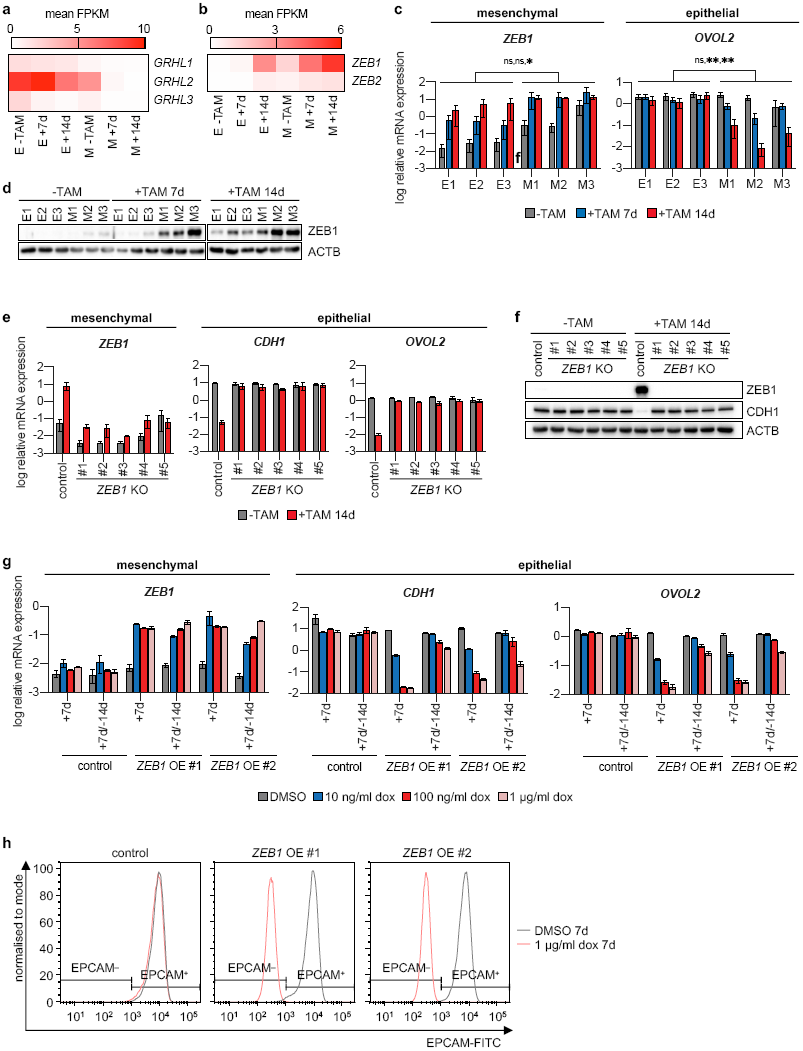
The EMT-transcription factor ZEB1 is required for EMT, but not sufficient to overcome EMT-resistance. **a**, Heatmap of mean FPKM values of RNA-sequencing data of *GRHL1*-*3* of 3 E-SCCs (E) and 3 M-SCCs (M), untreated (-TAM), treated with Tamoxifen for 7 days (+7d), or treated with Tamoxifen for 14 days (+14d). **b**, Heatmap of mean FPKM values of RNA-sequencing data of *ZEB1*-*2* of 3 E-SCCs (E) and 3 M-SCCs (M), treated as described in a. **c**, log relative mRNA expression levels of *ZEB1* and *OVOL2* in E-SCCs (E1-E3) and M-SCCs (M1-M3), treated as described in a. n=3; mean ± SEM; multiple t-tests (Holm-Sidak correction); p-values: *<0.05, **<0.005, ns=not significant. **d**, Immunoblot of ZEB1 and β-actin (ACTB) of E-SCCs (E1-E3) and M-SCCs (M1-M3), treated as described in a. **e**, log relative mRNA expression levels of *ZEB1, CDH1*, and *OVOL2* of a M-SCC control clone and M-SCC *ZEB1* knockout clones (*ZEB1* KO #1-5), untreated (-TAM) or treated with Tamoxifen for 14 days (+TAM 14d). n=2; mean ± SEM. **f**, Immunoblot of ZEB1, E-cadherin (CDH1), and β-actin (ACTB) of a M-SCC control clone and M-SCC *ZEB1* knockout clones (*ZEB1* KO #1-5) treated as described in e. **g**, log relative mRNA expression levels of *ZEB1, CDH1*, and *OVOL2* of an E-SCC control clone and E-SCC *ZEB1* overexpression clones (*ZEB1* OE #1, #2) treated with DMSO or with 10 ng/ml, 100 ng/ml, or 1 µg/ml doxycycline (dox) for 7 days (+7d), or after 7 days of treatment followed by 14 days of withdrawal (+7d/-14d). n=3 technical replicates; mean ± SEM. **h**, Flow cytometric staining of EPCAM of an E-SCC control clone and E-SCC *ZEB1* overexpression clones (*ZEB1* OE #1, #2) treated with DMSO or with 1 µg/ml doxycycline (dox) for 7 days (7d).

Together these results suggest that differences in ZEB1-activation and -levels are implicated in the contrasting responses to EMT-induction in E-SCCs and M-SCCs. In detail, Twist1-activation resulted in higher *ZEB1* expression in EMT-susceptible cells, which in turn lead to the transcriptional repression of epithelial genes such as *EPCAM, CDH1, OVOL2*, and *MIR200*-family members. By contrast, E-SCCs resisted repression of epithelial genes, which also served to restrict ZEB1-activation since many of these genes are known to function in a direct negative feedback loop.

To test, whether *ZEB1* was necessary for the irreversible EMT observed in M-SCCs, we generated M-SCC *ZEB1* knockout (KO) single-cell clones (Figure 5e, f, Supplementary Figure 5b, c). Upon Twist1-activation through TAM treatment, *ZEB1* KO clones indeed failed to undergo EMT in contrast to control cells, as evidenced by the maintenance of downstream epithelial markers like *CDH1* and *OVOL2* (Figure 5e, f, Supplementary Figure 5d).

These results demonstrate that activation of *ZEB1* is required for EMT and the resultant trans-differentiation from an epithelial to a mesenchymal phenotype alongside with the repression of epithelial markers. Therefore, inefficient induction of *ZEB1* expression in E-SCCs might explain the observed resistance to EMT-induction. Next, we wished to determine whether *ZEB1* overexpression (OE) was sufficient to overcome EMT-resistance in E-SCCs. Therefore, we generated E-SCC single-cell clones conditionally overexpressing *ZEB1*. Treatment with doxycycline and concomitant induction of *ZEB1* expression for 7 days led to EMT in *ZEB1* OE clones as indicated by morphology and downregulation of its targets *OVOL2, CDH1*, and EPCAM in contrast to a control clone, which remained resistant to EMT (Figure 5g, h, Supplementary Figure 5e, f). However, upon doxycycline withdrawal for 14 days, *ZEB1* OE clones reverted to an epithelial phenotype and regained expression of epithelial markers such as *CDH1* and *OVOL2* (Figure 5g, Supplementary Figure 5e).

Thus, even though we observed a transient downregulation of epithelial traits in E-SCCs, *ZEB1* overexpression was not sufficient for trans-differentiation of EMT-resistant cells into an irreversible mesenchymal state.

Together, these data suggest that a globally acting epigenetic mechanism exists in EMT-resistant cells that prevents ZEB1 from repressing epithelial genes independently of *ZEB1* expression levels. The resultant epigenetic stabilization of cellular identity promotes plasticity by enabling cells to maintain an epithelial identity while co-expressing mesenchymal markers. Moreover, it appears that initiation of proliferation in different settings that mimic metastatic outgrowth is directly connected to epithelial identity, whereas this capacity is suppressed by loss of epithelial traits through mesenchymal reprogramming.

## Discussion

In this study, we report that heterogeneity exists with respect to EPCAM levels which range from low to high in patient-derived metastatic breast cancer samples obtained from liquid biopsies. We observed that the ability to initiate tumours upon xenotransplantation was enriched in EPCAM^high^ compared to EPCAM^low^ patient-derived metastatic breast cancer cells. The transcriptome of EPCAM^low^ breast cancer cells showed a correlation with EMT related genes, suggesting that the maintenance of an epithelial identity in EPCAM^high^ breast cancer cells promotes metastatic outgrowth of these cells (Figure 6a). Indeed, using an experimental system based on human breast cancer cells, we observed that some cells actively resist EMT-induction. This resistance corresponded to a global epigenetic mechanism serving to restrict the activity of the transcriptional repressor ZEB1 and preventing the loss of chromatin accessibility at loci containing epithelial genes including E-cadherin (*CDH1*, Figure 6b). Indeed, expression of E-cadherin has recently been shown to directly promote metastatic outgrowth^27^.

**Figure 6:**
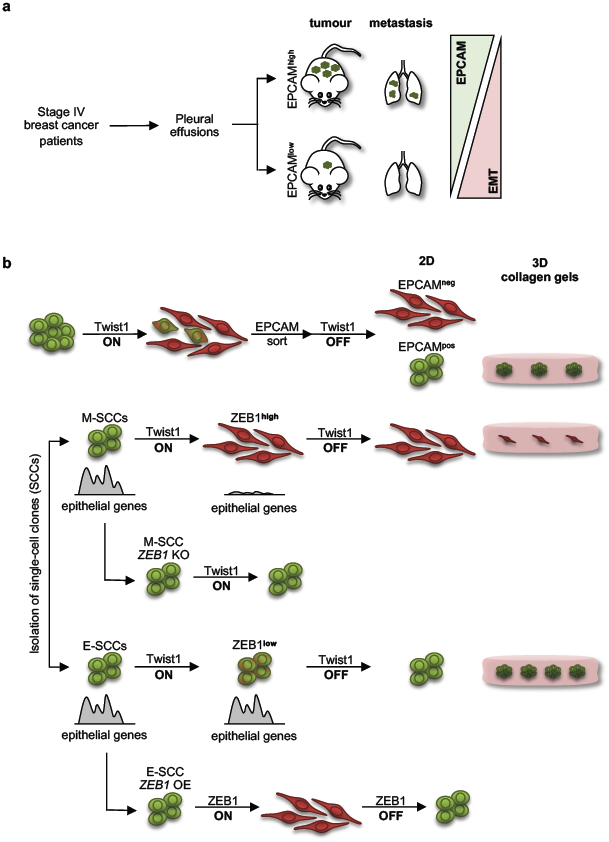
Maintenance of epithelial features promotes metastatic outgrowth. **a**, High EPCAM expression of metastatic breast cancer cells derived from pleural effusions of stage IV breast cancer patients enriches for tumour initiation and progression (EPCAM^high^). On the contrary, reduced EPCAM levels of metastatic breast cancer cells prevent tumour initiation and progression and correlate with EMT (EPCAM^low^). **b**, Upon Twist1-activation of HMLE-Twist1-ER cells, a subset of cells expresses high levels of ZEB1, represses the expression of epithelial genes, and undergoes stable and irreversible EMT, leading to loss of colonizing capacity in a three-dimensional environment (EPCAM^neg^/M-SCCs). Knockout (KO) of ZEB1 in these cells inhibits trans-differentiation into a mesenchymal phenotype upon an EMT-stimulus. Another subset of cells expresses low levels of ZEB1 and resists repression of epithelial genes, resulting in resistance to EMT and maintenance of the colonizing capacity (EPCAM^pos^/E-SCCs). Transient overexpression (OE) of ZEB1 within these cells activates EMT temporarily, but does not allow stable and irreversible trans-differentiation to a mesenchymal state.

Importantly, our data not only supports the concept that epithelial features promote metastatic outgrowth, but reveal that these features are actively maintained. On the flip side, we show that breast cancer cells that are susceptible to repression of epithelial features undergo irreversible EMT, a process characterized by high levels of ZEB1 and concomitantly, reduced chromatin accessibility over widespread genomic regions that include genes determining epithelial identity and, at the same time restricting ZEB1-activity via negative feedback loops (Figure 6b). When such cells with compromised epithelial identity are either derived from breast cancer metastatic biopsies or cell lines, we consistently observe a reduced ability to initiate proliferation in different settings that mimic metastatic outgrowth.

In conclusion, our study combines evidence from multiple models of the human disease, consistently indicating that the preservation of epithelial gene expression during EMT-induction is a fundamental mechanism for metastatic propagation of breast cancers, whereas EMT-susceptibility results in a permanent mesenchymal state coupled to growth arrest.

Our findings are supported by a recent study showing that spontaneous epithelial-mesenchymal trans-differentiation in a subset of HMLE-Ras cells residing in a mesenchymal state reduces tumorigenicity^28^. This study, together with a comprehensive analysis of EMT-transition states in murine models of mammary and cutaneous squamous carcinoma also suggests that a hybrid epithelial-mesenchymal phenotype is required for tumour progression^11,28^. However, neither the work by Padmanaban et al.^27^ nor our study is necessarily in conflict with these data. On the contrary, our work suggests a possible mechanistic basis for a hybrid epithelial-mesenchymal state, defined by active maintenance of epithelial gene expression and concomitant upregulation of mesenchymal markers. Moreover, our data suggest that transient entrance into such a hybrid state might enable metastatic progression, which is in line with several other studies^4,6,7^.

However, when focusing on the last step of metastasis, i.e. the outgrowth of disseminated cells into actively growing macrometastases, our data support a model suggesting that epithelial features, not necessarily a hybrid phenotype, drive metastatic outgrowth, as epithelial identity appears to be tied to cellular programs that govern proliferation. This is particularly important in light of the fact that it is this last step of metastatic progression that limits patient survival. In line with that, others hypothesized that repression of EMT will not affect outgrowth of metastases, as epithelial tumour cells with high proliferative capacity outcompete cells that have undergone an EMT^9^. One obvious question arising from our data is why EPCAM^low^ breast cancer cells are present in metastatic effusion biopsies in the first place, if they are not capable of propagating the disease. In this context, it has been previously observed that in primary tumours, mesenchymal cancer cells might cooperate with epithelial cancer cells to enable the systemic dissemination of both cell populations^29^. Therefore, one could hypothesize that EPCAM^low^ cells might promote the systemic dissemination and subsequent growth of EPCAM^high^ breast cancer cells.

Additionally, we need to keep in mind that patients within this study were treated with chemotherapy. As it was shown that mesenchymal, EMT cells resist chemotherapy whereas epithelial, non-EMT tumour cells are sensitive to chemotherapy^9^, the presence of mesenchymal breast cancer cells at the metastatic site could also be explained by their resistance to the therapy. These cells could potentially remain as dormant population and thereby counterbalance their proliferative disadvantage.

Because of that, we would like to add an important caveat to these considerations: our data suggest that EMT might create heterogeneity with respect to proliferative capacity. Although our data suggest that complete EMT, and hence, the resultant growth suppression might be permanent, it will nonetheless be important to take the observed cellular heterogeneity into account when developing new treatment strategies for metastatic breast cancer.

Most importantly, our results strongly support the emerging evidence linking epithelial identity to metastatic outgrowth. Additionally, our data provide evidence for a globally acting, epigenetic mechanism serving to render cells resistant against mesenchymal reprogramming and thereby serve to actively maintain and anchor epithelial features in the metastases.

## Methods

### Clinical specimens

Pleural and ascitic effusions were obtained from post-menopausal stage IV patients admitted to the Division of Gynecologic Oncology, National Centre for Tumor Diseases and the Women’s Clinic of the Mannheim University Hospital. The study was approved by the ethical committee of the University of Heidelberg (case number S295/2009) and University of Mannheim (2010-024238-46). All samples were transferred to the laboratory no later than 1 hour after thoracentesis or laparocentesis and processed to extract cancer cells as previously described^30^. Patient’s diagnostic information and clinical disease characteristics are described in detail in Table 1.

### Xenograft assay for patient-derived metastatic breast cancer cells

Female NOD.Cg-Prkdc^scid^ Il2rg^tm1Wjl^/SzJ (NSG) mice were transplanted at 6 to 8 weeks of age. A suspension of FACS-purified cells in sterile PBS was combined to growth factor-reduced Matrigel (BD) in a 1:1 ratio and then injected in the 4^th^ or the 5^th^ mammary fat pad. All mice received subcutaneous implantation of beta-estradiol as solid pellets (Innovative Research of America) with modalities that were already described^2^. Engraftment of cancer cells and growth *in vivo* was monitored by regular palpation of the implantation site. Animal care and procedures followed the German legal regulations and were previously approved by the governmental review board of the state of Baden-Württemberg, operated by the local Animal Welfare Office (Regierungspräsidium Karlsruhe) under the license number G240-11.

### Immunohistochemical analysis of tumour xenografts

Tumour lesions generated in mice and metastatic lungs were freshly collected upon necropsy and embedded in paraffin for downstream histological inspections. Sections were obtained, processed and stained as described^31^. For immunohistochemistry, the following antibodies were used: anti-human Ki-67 (DAKO, clone Ki-67), anti-oestrogen receptor alpha (Thermo Fisher Scientific, clone SP1), anti-progesterone receptor (DAKO, clone PgR636), anti-HER2 (DAKO, polyclonal serum A0485), anti-CDH1 (DAKO, clone M3612), and anti-Vimentin (DAKO, clone M0725).

### Targeted next-generation sequencing of somatic mutations

Genomic DNA was extracted from FACS-purified cancer cells and matched germline controls (CD45^+^ white blood cells purified by FACS sorting from the same metastatic effusion). The concentration of nucleic acids was assessed by fluorimetric measurement through QuBit 3.0 and the amount of amplifiable DNA (sequencing-grade quality) was determined using a quantitative assay (TaqMan RNAseP detection assay) on a StepOnePlus instrument (both: Thermo Fisher Scientific). Samples were amplified using a custom-designed gene panel for breast cancer^32^ covering the most recurrent mutations^33^. Library preparation and sequencing were performed using the multiplex PCR-based Ion Torrent AmpliSeq™ technology (Thermo Fisher Scientific) and Ion S5XL technology, with procedures already described^34^.

### RNA-sequencing of patient-derived metastatic breast cancer cells

RNA extraction was carried out from 40,000 FACS-sorted cells using ARCTURUS PicoPure RNA isolation kit (Thermo Scientific, Cat. KIT0204). RNA quality was assessed using BioAnalyzer 2100 (Agilent). Libraries were prepared from 1 μg of total RNA using a TruSeq Stranded Total RNA Kit with Ribo-Zero Human/Mouse/Rat (Illumina) according to the manufacturer’s instructions. The pooled RNA libraries were sequenced on an Illumina HiSeq2000 to obtain 100-bp paired-end reads. RNA-sequencing reads were aligned to the human reference genome hg19 and quantified using the stranded option of STAR version 2.4.2a30. Furthermore, the DESeq2 package was used to obtain normalized expression values as described^35^. The GSEA desktop application was utilized for enrichment analysis of the indicated gene sets. Gene set databases downloaded from the Broad Institute website or custom-derived signatures were analyzed using default settings with 2000 permutations.

### Cell lines

HMLE-Twist1-ER, generated as previously described^36,37^, were kindly gifted by Robert A. Weinberg (Whitehead Institute). HMLE-Twist1-ER (CD24^high^) cells were cultured in Mammary Epithelial Cell Basal Medium containing 0.004 ml/ml BPE, 10 ng/ml EGF, 5 μg/ml Insulin and 0.5 μg/ml hydrocortisone (PromoCell) supplemented with Pen/Strep (Invitrogen) and Blasticidin S HCl (Thermo Fisher Scientific) at a final concentration of 10 µg/ml. To obtain HMLE-Twist1-ER CD24^high^ single-cell clones (SCCs), cell were seeded to a 96-well plate (0.3 cells/well). Wells were checked by eye for single-cells. Wells with a single cell were further passaged and expanded. For Twist1-activation, cells were treated with (Z)-4-Hydroxytamoxifen (TAM, Sigma-Aldrich) at a final concentration of 20 nM for the indicated number of days. HMLE-Twist1-ER *ZEB1* overexpression cells were isolated with cloning cylinders (Sigma) and propagated in medium further supplemented with Puromycin Dihydrochloride (Thermo Fisher Scientific) at a final concentration of 1 µg/ml. For tracing experiments, E-SCC cells were lentivirally transduced with pRRL-cPPT-CMV-GFP-W and fluorescence-activated cell sorting (FACS) was used to select GFP-positive cells. SUM-149PT cells were cultivated in F12 containing 5% FBS. SUM-149PT cells were authenticated using Multiplex Cell Authentication (Multiplexion, Heidelberg, Germany) as described^38^. The SNP profiles matched known profiles or were unique. The purity of SUM-149PT cells was validated using the Multiplex cell Contamination Test (Multiplexion, Heidelberg) as described^39^. The medium of all cells was changed every two to three days.

### RNA isolation and qRT-PCR

RNA was isolated using the miRNeasy Mini Kit (Qiagen) or the mRNeasy Mini Kit (Qiagen). Reverse transcription of mRNA was performed with the EasyScriptPlus Kit. For qRT-PCR, the Power SYBR Green-PCR Master Mix (Applied Biosystems) was used. *RPL32* was used as a control for normalization. Reverse transcription of miRNAs was performed with the miScript RT Kit (Qiagen) and qRT-PCR was performed with the Power SYBR Green-PCR Master Mix (Applied Biosystems) and miScript Primer Assays (control/normalization:HS-RNU6-2_11, *MIR141*:HS-miR-141_1, *MIR200A*:HS-miR-200a_1, *MIR200B*:HS-miR-200b_3, *MIR200C*:HSmiR200c_1, Qiagen). Samples ran on a QuantStudio 12K Flex qPCR system (Life Technologies). Data were analysed with the ΔCt method and relative expression levels are displayed as 1000 × 2^-ΔCt^. Following primer pairs were used: *CDH1*: forward 5’-TGCCCAGAAAATGAAAAAGG-3’ and reverse 5’- GTGTATGTGGCAATGCGTTC-3’, *EPCAM*: forward 5’- ATAACCTGCTCTGAGCGAGTG-3’ and reverse 5’- TGCAGTCCGCAAACTTTTACTA-3’, *FN1*: forward 5’- CAGTGGGAGACCTCGAGAAG-3’ and reverse 5’- TCCCTCGGAACATCAGAAAC-3’, *Twist1*: forward 5’-GGACAAGCTGAGCAAGATTCA-3’ and reverse 5’- CGGAGAAGGCGTAGCTGAG-3’, *OVOL2*: forward 5’- ACAGGCATTCGTCCCTACAAA-3’ and reverse 5’- CGCTGCTTATAGGCATACTGC- 3’, *RPL32*: forward 5’- CAGGGTTCGTAGAAGATTCAAGGG-3’ and reverse 5- ‘CTTGGAGGAAACATTGTGAGCGATC-3’, *VIM*: forward 5’- GAGAACTTTGCCGTTGAAGC-3’ and reverse 5’- GCTTCCTGTAGGTGGCAATC-3’, *WNT5A*: forward 5’- ATGGCTGGAAGTGCAATGTCT-3’ and reverse 5’- ATACCTAGCGACCACCAAGAA-3’, *ZEB1*: forward 5’- GCACAAGAAGAGCCACAAGTAG-3’ and reverse 5’- GCAAGACAAGTTCAAGGGTTC-3’.

### Immunofluorescence

Cells were cultured on coverslips coated with poly-D-lysine (Sigma-Aldrich), in floating 3D-collagen gels, or on murine lung slices. Cells were fixed with 4% paraformaldehyde for 10 to 15 minutes and permeabilized with 0.2% Triton X-100. After blocking with 10% normal goat serum in 0.1% BSA (Sigma-Aldrich), coverslips, floating 3D-collagen gels, and murine lung slices were incubated overnight with primary antibodies in 0.1% BSA as follows: anti-CDH1 (clone EP700Y, Biozol, 1:250) or anti-CDH1-Alexa-488 (clone 24E10, New England Biolabs, 1:50), anti-Twist1 (clone Twist2C1a, Santa Cruz, 1:500), anti-VIM (clone V9, Abnova, 1:100). Secondary antibodies were coupled to Alexa-488, −594, or −546 (Life Technologies, 1:250). Cell nuclei were visualized with 40,6-diamidino-2-phenylindole-dihydrochloride (DAPI, Sigma-Aldrich). Slides were mounted with Aqua-Poly/Mount reagent (Polysciences). Corrected total cell fluorescence (CTCF) of DAPI was calculated using the following formula: CTCF = integrated density – (area of selected cells x mean fluorescence of background readings). Pictures were taken with a Zeiss Axio Imager.M2 using Zen software or with an Olympus Confocal using FV10-ASW software.

### Fluorescence-Activated Cell Sorting (FACS) and Flow Cytometry

Metastatic breast cancer cells were stained in a 1% BSA solution containing 2 mM Ethylenediaminetetraacetic acid (EDTA) under the presence of FcR blocking reagent (Miltenyi). HMLE-Twist1-ER cell suspensions were stained in Mammary Epithelial Cell Basal Medium (PromoCell). Propidium Iodide (PI), or 7-AAD (BD Biosciences, 1:125 were used to exclude dead cells, depending on the staining panel. Cells were sorted and analysed on a FACSAria Illu, an LSR Fortessa, and a FACS Aria Fusion (all BD) and further analysed with FlowJo. Antibodies: anti-EPCAM-FITC (VU-1D9 clone, Biozol, 1:25, or REA764 clone, Miltenyi, 1:11), anti-CD45-VioBlue (REA747 clone, Miltenyi, 1:11).

### Immunoblotting

Adherent cells were harvested by scraping on ice after 1 wash with PBS. Collected cell pellets were resuspended in RIPA buffer (50 mM Tris pH 8.0, 150 mM NaCl, 1% NP-40, 0.5% sodium deoxycholate, 0.1% sodium dodecyl sulphate and 5 mM EDTA pH 8.0) containing a protease and a phosphatase inhibitor (Sigma-Aldrich). Protein concentration was measured using the DC Protein Assay (Bio-Rad Laboratories). 10 µg protein were separated on 10% SDS-PAGE gels and transferred on PVDF membranes by wet-blot transfer. Immunoblotting was performed using the following primary antibodies: CDH1 (clone EP700Y, Biozol, 1:25000), ZEB1 (clone H-102, Santa Cruz Biotechnology, 1:200) or ZEB1 (Sigma, 1:5000), ACTB (clone AC-15, Abcam, 1:6000) was used as a loading control. Secondary antibodies (anti-mouse and anti-rabbit) were conjugated to horseradish peroxidase (Jackson Immuno Research, 1:12500). Chemiluminescence reaction was activated with ECL Prime Western Blotting Detection Reagent (GE Healthcare) and detection was carried out on a ChemiDoc Imaging System using Image Lab software (Bio-Rad Laboratories).

### Floating 3D-collagen gels

Floating 3D-collagen gels with a final concentration of 1.3 mg/ml collagen I (collagen type I from rat tail, Corning) were prepared as previously described^7,15^. Briefly, 300 HMLE-Twist1-ER cells were embedded per collagen gel. Cells were further cultivated in Mammary Epithelial Cell Basal Medium containing 0.004 ml/ml BPE, 10 ng/ml EGF, 5 μg/ml Insulin and 0.5 μg/ml hydrocortisone (PromoCell) supplemented with Pen/Strep (Invitrogen) and Blasticidin S HCl (Thermo Fisher Scientific) at a final concentration of 10 µg/ml. Medium was changed every two to three days. After 10 days in culture, floating 3D-collagen gels were fixed with 4% paraformaldehyde for 15 minutes. Immunofluorescent staining was performed as described above. To visualize colonies, floating 3D-collagen gels were incubated in a carmine alum staining solution (2 g/l carmine (Sigma-Aldrich), 5 g/l aluminium potassium sulphate (Sigma-Aldrich), two crystals of thymol) at 4°C overnight. Pictures of gels were taken with a Leica DFC450 C Stereomicroscope with a Plan 0.8x LWD objective and the Leica Application Suite V4 software. Pictures were stitched with Panorama Stitcher. Carmine positive colonies were counted using ImageJ.

### Colonizing assay on murine precision cut lung slices (PCLS)

PCLS were generated as previously described^17^. Briefly, C57BL6/N mice of 8-12 weeks were intubated and after the dissection of the diaphragm, lungs were flushed through the heart with sterile sodium chloride solution. Using a syringe pump, lungs were filled with low gelling temperature agarose (2% by weight, A9414; Sigma; kept at 40°C) in sterile cultivation medium (DMEM/Ham’s F12 (Gibco) supplemented with 100 U/ml penicillin, 100 μg/ml streptomycin and 2.5 μg/ml amphotericin B (Sigma)). Separated lobes were cut with a vibratome (Hyrax V55; Zeiss, Jena, Germany) to a thickness of 300 μm using a speed of 10–12 μm/s, a frequency of 80 Hz and an amplitude of 1 mm. The PCLS were cultivated in sterile cultivation medium containing 0.1%FCS. HMLE-Twist1-ER cells were lentivirally transduced with pRRL-cPPT-CMV-GFP-W. Fluorescence-activated cell sorting (FACS) was used to select GFP positive cells. 2×10^5^ cells were plated per lung slice and cultured in Mammary Epithelial Cell Basal Medium containing 0.004 ml/ml BPE, 10 ng/ml EGF, 5 μg/ml Insulin and 0.5 μg/ml hydrocortisone (PromoCell) supplemented with Pen/Strep (Invitrogen) for 5 days, changing the medium every day.

### CRISPR/Cas9 of *ZEB1*

We made use of the transient expression of Cas9 and four guide RNAs (gRNAs) targeting two regions of exons that are found in all 9 transcript variants of *ZEB1*. Guide RNAs (gRNAs) were designed with Benchling and the MIT CRISPR design tool (http://crispr.mit.edu/). For creating a vector harbouring all four *ZEB1* gRNAs, gRNAs were cloned by applying the string assembly gRNA cloning (STAgR) protocol^40^. For Gibson Assembly, a Gibson Assembly Master Mix (New England Biolabs) was used according to manufacturer’s instructions. For transformation into XL10-Gold ultracompetent cells (Agilent Technologies) the Gibson Assembly was diluted 1:4. Purified plasmids were transfected into HMLE-Twist1-ER cells. In detail, 24h before transfection, 1.5 – 2×10^5^ cells were seeded in 6-well dishes. Cells were then transfected with a Cas9-GFP-expressing plasmid (pSpCas9(BB)-2A-GFP) and the gRNA containing STAgR-Neo plasmid using the TransIT-X2 transfection reagent (Mirus Bio LLC) according to manufacturer’s instructions. As a control, cells were transfected with the Cas9-GFP plasmid and a plasmid harbouring the guide RNA scaffold and termination sequence. After 2.5 days cells were sorted for GFP expression on a FACSAriaIIIu (BD Biosciences). To obtain single-cell clones, sorted cells were seeded into 96-well plates (1 cell/well). Wells were checked by eye for single-cells. Wells with a single-cell were further passaged and expanded. To screen for cells with successful deletion, cells were lysed in DNAzol reagent (Thermo Fisher Scientific) and genomic DNA (gDNA) of *ZEB1* knockout single-cell clones was isolated with the DNAzol Reagent (Invitrogen) according to the manufacturer’s instructions for adherent cells (monolayer). PCRs of the specific loci were performed to screen for DNA modifications and *ZEB1* knockout was validated by immunoblotting.

Following gRNAs were used: *ZEB1* NM128128 Exon 5: 5’- GCCTCTATCACAATATGGAC-3’ and 5’-ACAACTCAGCCCTCAATGGA-3’, *ZEB1* NM128128 Exon 7: 5’-AGTTCTGTCACAAGCATGCA-3’ and 5’- TTGCCGTATCTGTGGTCGTG-3’.

Following PCR primers were used: *ZEB1* NM128128 Exon 5: forward 5’- GCATAGGGACTCAGTGGAAACT-3’ and reverse 5’- AGGAGGCAACTCCCTTTACTAC-3’, *ZEB1* NM128128 Exon 7: forward 5’- GGTCGGTGAAATGGGATAAGAAAAA-3’ and reverse 5’- ACCACCAGTGAAAACCCCATT-3’.

### Overexpression of *ZEB1*

To overexpress *ZEB1*, we made use of the All-in-One piggyBac system designed for inducible transgene expression^41^. In detail, the *ZEB1* (NM1174096) cDNA was cloned into the pENTR1A-no ccdb vector (#17398, Addgene) to generate the pENTR1A-*ZEB1* vector. Next, the Gateway LR Clonase II Enzyme mix (Invitrogen) was used to transfer the *ZEB1* cDNA from the pENTR1A-*ZEB1* vector into the PB-TAC-ERP2 (#80478, Addgene) to generate the PB-*ZEB1* vector. Cells were transfected with the linearized PB-ZEB1 plasmid and the pCMV-hyPBase vector (kind gift of Roland Rad, TU Munich) in a ratio of 4:1, respectively, using the TransIT-X2 transfection reagent (Mirus Bio LLC) according to manufacturer’s instructions. For selection, cells were cultured in medium containing 1 µg/ml Puromycin Dihydrochloride (Thermo Fisher Scientific) 48 hours after transfection. Next, single-cell colonies were isolated using cloning cylinders (Sigma) and expanded for further analysis.

### RNA-sequencing of HMLE-Twist1-ER cells

Library preparation was performed using the TruSeq Stranded Total RNA Library Prep Kit with Ribo-Zero (Illumina). Briefly, total RNA was isolated using the miRNeasy Mini Kit (Qiagen) and RNA integrity number (RIN) was determined with the Agilent 2100 BioAnalyzer (RNA 6000 Nano Kit, Agilent). For library preparation, 1 μg of RNA was depleted for cytoplasmatic rRNAs, fragmented, and reverse transcribed with the Elute, Prime, Fragment Mix. A-tailing, adaptor ligation, and library enrichment were performed as described in the High Throughput protocol of the TruSeq RNA Sample Prep Guide (Illumina). RNA libraries were assessed for quality and quantity with the Agilent 2100 BioAnalyzer and the Quant-iT PicoGreen dsDNA Assay Kit (Life Technologies). RNA libraries were sequenced as 150 bp paired-end runs on an Illumina HiSeq4000 platform. The STAR aligner (v 2.4.2a) with modified parameter settings (--twopassMode=Basic) is used for split-read alignment against the human genome assembly hg19 (GRCh37) and UCSC knownGene annotation^42^. To quantify the number of reads mapping to annotated genes we use HTseq-count (v0.6.0)^43^. FPKM (Fragments Per Kilobase of transcript per Million fragments mapped) values are calculated using custom scripts. PCA plots were created with the R package ggplot2^44^.

### Omni-ATAC-sequencing of HMLE-Twist1-ER cells

ATAC-sequencing was done as described previously^45^. Briefly, 50,000 cells (viability > 90%) were harvested by trypsinisation and pelleted (500 rcf, 4°C, 5 min). If the cell viability of cell suspensions used for ATAC-seq was below 90%, dead cells were removed with the Dead Cell Removal Kit (Milteny Biotech) according to the manufacturer’s instructions and then harvested as the others. Cell pellets were resuspended in ATAC Resuspension Buffer (10mM Tris-HCl pH 7.4, 10 mM NaCl, 3 mM MgCl_2_) supplemented with 0.1% NP40, 0.1% Tween-20 and 0.01% digitonin for lysis, incubated on ice for 3 min and then 1 ml of ATAC Resuspension Buffer supplemented only with 0.1% Tween-20 was added and spun (500 rcf, 4°C, 10 min) to collect nuclei. Nuclei were subsequently re-suspended in 50 µl of Transposase Reaction containing 25 µl 2x Tagmentation Buffer (20 mM Tris-HCl pH 7.6, 10 mM MgCl_2_, 20% dimethyl Formamide, H_2_O), 2.5 µl Tn5 Transposase (Illumina Nextera DNA Library Preparation Kit), 5.25 µl H_2_O, 16.5 µl PBS, 0.25 µl of 2% digitonin (Promega) and 0.5 µl of 10% Tween-20. Reactions were incubated for 30 min at 37°C in a thermomixer shaking at 900 rpm and DNA was then purified using Qiagen PCR clean-up MinElute kit (Qiagen). Transposed DNA was subsequently amplified in 50 µl reactions with custom primers as described previously^46^. After 4 cycles amplification, libraries were then monitored with qPCR: 5 µl PCR sample in a 15 µl reaction with the same primers. qPCR output was monitored for the ΔRN; 0.25 ΔRN cycle number was used to estimate the number of additional cycles of the PCR reaction needed for the remaining PCR samples. Amplified libraries were purified with the Qiagen PCR clean-up MinElute kit and size selected for fragments less than 600 bp using the Agencourt AMPure XP beads (Beckman Coulter). Libraries were quality controlled by Qubit and Agilent DNA Bioanalyzer analysis. Deep sequencing was performed on a HiSeq 1500 system according to the standard Illumina protocol for 50bp single-end reads. ATAC-seq reads were aligned to the human genome hg38 using Bowtie^47^ with options “-q –n 2 --best --chunkmbs 2000 -p 32 -S”. ATAC peaks over Input background were identified using Homer^48^ findPeaks.pl with option “-style factor”. Peaks from all samples were merged using mergePeaks resulting in a unified Peak set. The peak list was filtered for promoter-associated peaks (distance to TSS < 1000bp) with bedtools. Raw ATAC coverage counts were then calculated with annotatePeaks. PCA analysis was performed on ATAC peaks coverage data with the R function prcomp. The list of differential ATAC peaks was determined with the DESeq2 results function for all possible comparisons between the samples and filtered for adjusted p-value < 0.05 and log2 fold change > 3. The cluster heatmap was generated based on normalized coverage data of all differential ATAC peaks using the R pheatmap package in k_means mode with 12 clusters. Cluster-specific peaks were further analysed for transcription factor motifs using homer findMotifsGenome function in *de novo* mode.

### Data presentation and statistical analyses

Data are presented as mean±SEM of n=x experiments, with x indicating the number of independent experiments performed, unless otherwise stated. Statistical analysis was performed using GraphPad Prism software. In general, a p-value below 0.05 was considered significant.

## Data availability

The datasets that support the findings of this study have been deposited in the Gene Expression Omnibus (GEO) under the accession code GSE138329.

### Acknowledgements

We thank Christopher Breunig and Stefan Stricker (Helmholtz Center Munich) for sharing reagents and guidance for STAgR cloning, and Robert Weinberg (Whitehead Institute) for providing HMLE-Twist1-ER cells. We thank all members of the Scheel group, especially Lisa Meixner and Elena Panzilius, for sharing experimental expertise. We also thank Sandy Lösecke for technical assistance during RNA-sequencing. We thank Steffen Schmitt, Klaus Hexel, Ann Atzberger, and Marcus Eich from the DKFZ Flow Cytometry Core Facility for their kind assistance and all members of the DKFZ Laboratory Animal Core Facility for excellent animal welfare and husbandry. We are grateful to Roland Rad (Technical University Munich) for providing the hyperactive transposase for the PiggyBac assay. We further thank Heiko Lickert (Helmholtz Center Munich), Maximilian Reichert (Technical University Munich), Thordur Oskarsson, and Michael Milsom (HI-STEM Heidelberg) for constructive discussions and Nina Cabezas-Wallscheid (Max Planck Institute Freiburg) for her support. Finally, we thank Nicola Aceto (University of Basel) for helpful discussions and for his support. The work in the G.S. lab is funded by Deutsche Forschungsgemeinschaft (SFB1064 - A11 and SFB1321 - P13). This work was further supported by the Transluminal-B and Integrate-TN consortia funded by the Deutsche Krebshilfe, the Swiss-Bridge Award and the Dietmar Hopp Foundation (all to A.T.) and by a Max Eder Grant of the German Cancer Aid Foundation (Deutsche Krebshilfe 110225 to C.H.S.).

## Author Contributions

M.S., L.E., M.R.S., A.T., and C.H.S. conceived the project and designed experiments. L.E., M.S., H.D.M., C.K., J.M.B., M.R., V.V., S.H., and R.W. performed research and analysed the data. M.F., E.E., R.W., E.D., and S.H. analysed the data, provided crucial datasets and contributed to data interpretation. H.D.M. and G.S. performed ATAC-sequencing and data analysis. E.G. and T-M.S. performed RNA-sequencing. T.S. performed RNA-sequencing data analysis. M.L. and M.K. provided lung slices and the according assay. A.S., M.Sü., and S.S. provided patient samples, supervised and managed clinical data. N.P. and W.W. performed next-generation sequencing of mutational hotspots and supervised histopathological analysis of patient-derived xenografts. L.E. and M.S. assembled the figures. L.E., M.S., A.T. and C.H.S. wrote the manuscript. All authors read and approved the manuscript.

---

**Supplementary Figure 1:**
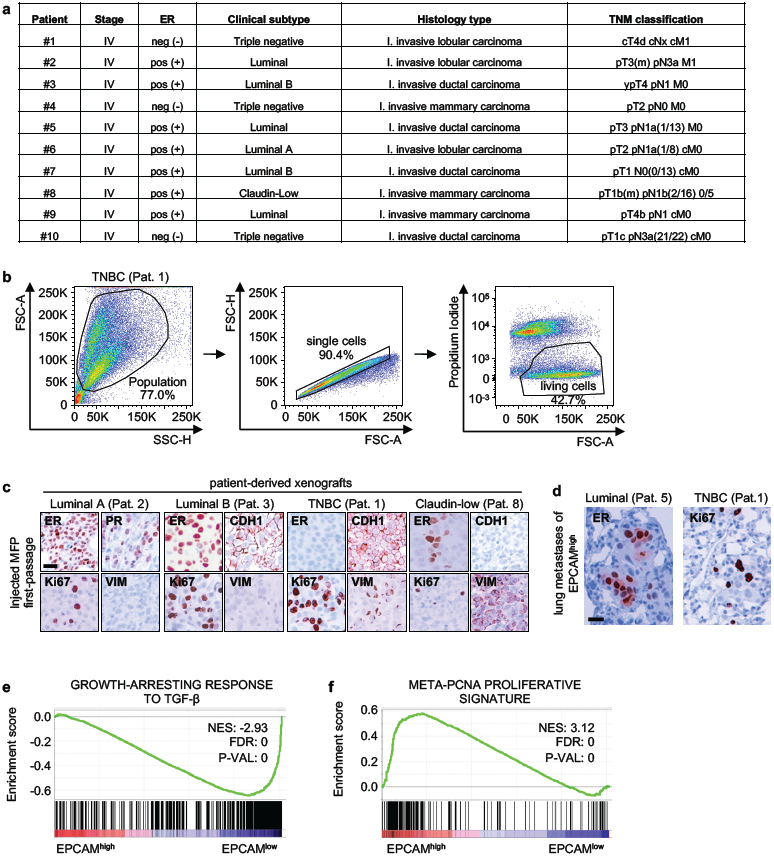
High EPCAM expression correlates with disease-propagating potential in metastatic breast cancer cells. **a**, Table showing tumour stage, oestrogen receptor (ER) status, clinical subtype, histology type, and TNM classification of clinical patients #1-10. **b**, Gating strategy for patient-derived metastatic breast cancer cells. TNBC, Triple Negative Breast Cancer; Pat., Patient. **c**, Immunohistochemical staining of oestrogen receptor (ER), progesterone receptor (PR), E-cadherin (CDH1), Ki67, and Vimentin (VIM) of tumour sections from engrafted mammary fat pads. The xenografts originate from cells of patients with the indicated breast cancer subtype. Scale bar: 25 µm. TNBC, Triple Negative Breast Cancer; Pat., Patient. **d**, Immunohistochemical staining of oestrogen receptor (ER) and Ki67 of spontaneous lung metastases originating from EPCAM^high^ cells of patients with the indicated breast cancer subtype. Scale bar: 25 µm. TNBC, Triple Negative Breast Cancer; Pat., Patient. **e**, Gene set enrichment analysis (GSEA) plot of the growth-arresting response to TGF-β of EPCAM^high^ versus EPCAM^low^ cells from an ER^+^ and an ER^−^ patient. NES, normalized enrichment score; FDR, false discovery rate; P-VAL, p-value. **f**, GSEA plot of the META-PCNA proliferative signature of EPCAM^high^ versus EPCAM^low^ cells from an ER^+^ and an ER^−^ patient. NES, normalized enrichment score; FDR, false discovery rate; P-VAL, p-value.

**Supplementary Figure 2:**
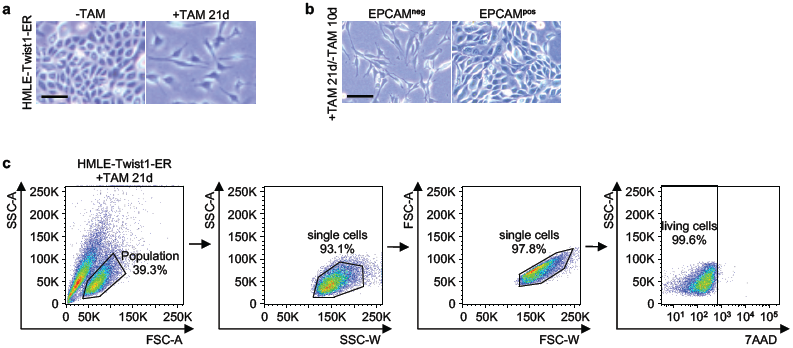
A subset of HMLE-Twist1-ER cells maintains EPCAM expression during EMT-induction. **a**, Bright field pictures of HMLE-Twist1-ER cells, untreated (-TAM) or treated with Tamoxifen for 21 days (+TAM 21d). Scale bar: 100 µm. **b**, Bright field pictures of sorted EPCAM^neg^ and EPCAM^pos^ HMLE-Twist1-ER cells after 10 days of Tamoxifen withdrawal (+TAM21d/-TAM10d). Scale bar: 100 µm. **c**, Gating strategy for HMLE-Twist1-ER cells.

**Supplementary Figure 3:**
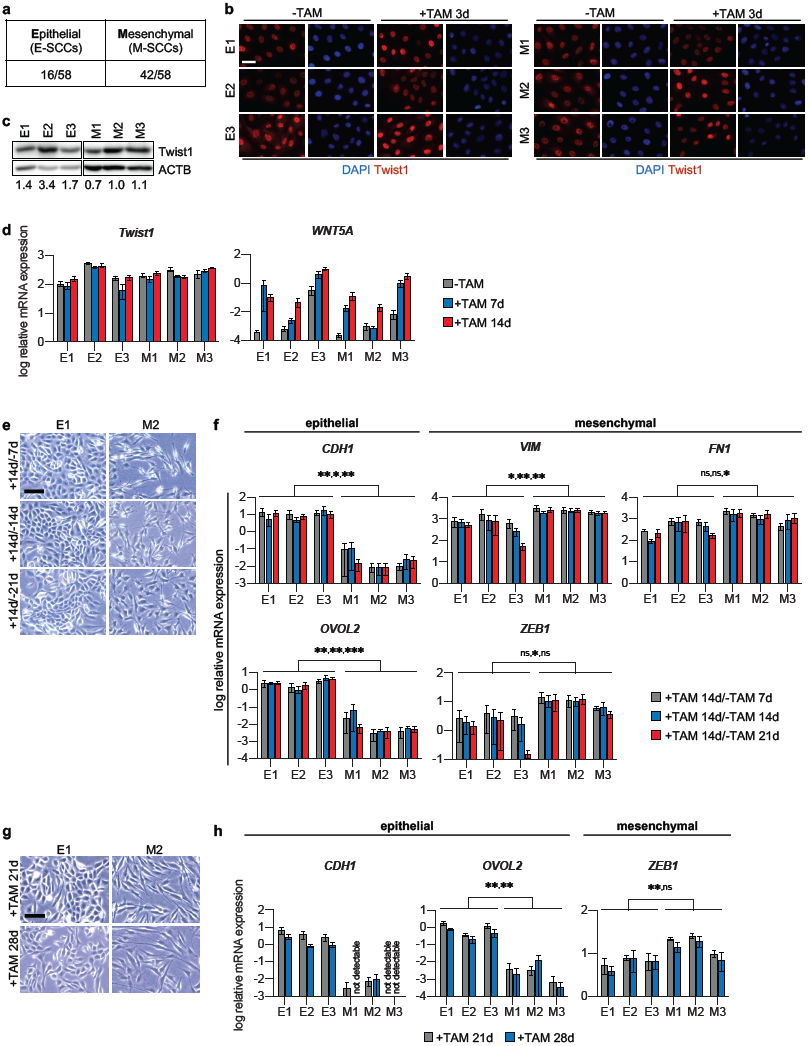
A subset of breast cancer single-cell clones resists complete EMT and maintains the ability to proliferate in different environments. **a**, Table showing number of epithelial (E-SCCs) and mesenchymal (M-SCCs) single-cell clones isolated from the HMLE-Twist1-ER bulk population and determined during Tamoxifen treatment. **b**, Immunofluorescence staining of DAPI (blue) and Twist1 (red) of E-SCCs (E1-E3) and M-SCCs (M1-M3), untreated (-TAM) or treated with Tamoxifen for 3 days (+TAM 3d), Scale bar: 20 µm. **c**, Immunoblot of Twist1 and β-actin (ACTB) of untreated E-SCCs and M-SCCs. Twist1 protein levels were quantified relatively to β-actin. **d**, log relative mRNA expression levels of *Twist1* and *WNT5A* in E-SCCs (E1-E3) and M-SCCs (M1-M3), untreated (-TAM), treated with Tamoxifen for 7 days (+TAM 7d), or treated with Tamoxifen for 14 days (+TAM 14d). n=3; mean ± SEM. **e**, Bright field images of a representative E-SCC (E1) and a representative M-SCC (M2), treated with Tamoxifen for 14 days followed by Tamoxifen withdrawal for 7 days (+14d/-7d), 14 days (+14d/-14d), or 21 days (+14d/-21d). Scale bar: 100 µm. **f**, log relative mRNA expression levels of *CDH1, VIM, FN1, OVOL2*, and *ZEB1* in E-SCCs (E1-E3) and M-SCCs (M1-M3), treated as described in e. n=3; mean ± SEM; multiple t-tests (Holm-Sidak correction); p-values: *<0.05, **<0.005, ***<0.0005, ns=not significant. **g**, Bright field images of a representative E-SCC (E1) and a representative M-SCC (M2), treated with Tamoxifen for 21 days (+TAM 21d) or for 28 days (+TAM 28d). Scale bar: 100 µm. **h**, log relative mRNA expression levels of *CDH1, OVOL2*, and *ZEB1* in E-SCCs (E1-E3) and M-SCCs (M1-M3), treated as described in g. n=3; mean ± SEM; multiple t-tests (Holm-Sidak correction); p-values: **<0.005, ns=not significant.

**Supplementary Figure 4:**
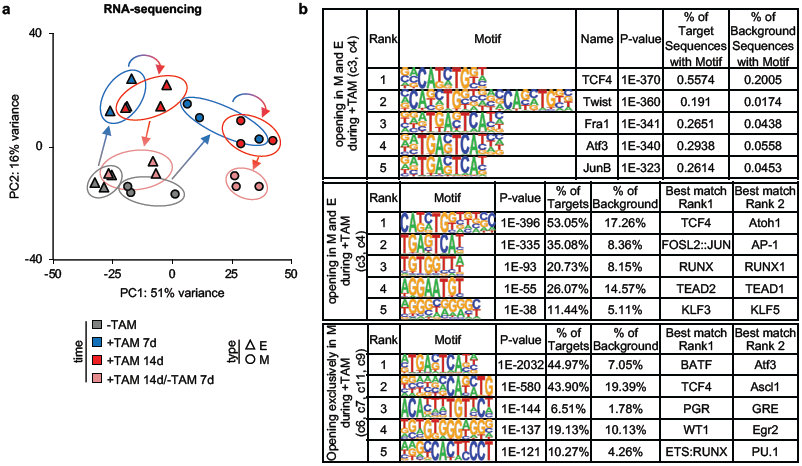
EMT-induction causes genome-wide chromatin and transcriptional changes. **a**, Principal component (PC) analysis of RNA-sequencing data of E-SCCs (△) and M-SCCs (**◯**), untreated (-TAM), treated with Tamoxifen for 7 days (+TAM 7d), treated with Tamoxifen for 14 days (+TAM 14d), or treated with Tamoxifen for 14 days followed by Tamoxifen withdrawal for 7 days (+TAM 14d/-TAM 7d). Each data point represents one SCC at the indicated time point. **b**, Top 5 hits of Homer *known motif* (upper table) or *de novo* (two lower tables) transcription factor motif analysis of grouped clusters opening in M-SCCs (M) and E-SCCs (E, two upper tables) or opening exclusively in M-SCCs (M, lower table) during Tamoxifen treatment (+TAM).

**Supplementary Figure 5:**
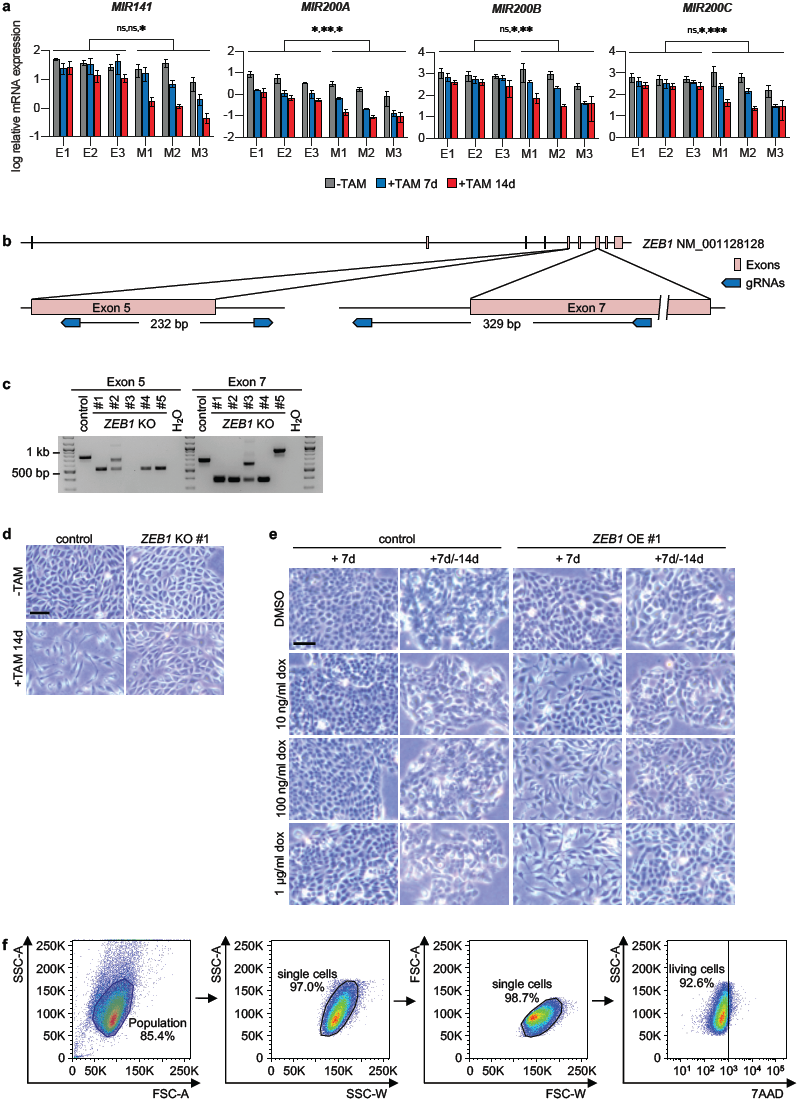
The EMT-transcription factor ZEB1 is required for EMT, but not sufficient to overcome EMT-resistance. **a**, log relative mRNA expression levels of *MIR141 and MIR200A*-*C* in E-SCCs (E1-E3) and M-SCCs (M1-M3), untreated (-TAM), treated with Tamoxifen for 7 days (+TAM 7d), or treated with Tamoxifen for 14 days (+TAM 14d). n=3; mean ± SEM; multiple t-tests (Holm-Sidak correction); p-values: *<0.05, **<0.005, ***<0.0005, ns=not significant. **b**, Schematic illustration of the localisation of guide RNAs (gRNAs) for CRISPR/Cas9 of the *ZEB1* (NM00128128) gene. **c**, PCR products of targeted DNA regions of a M-SCC control clone and M-SCC *ZEB1* knockout clones (*ZEB1* KO #1-5). **d**, Bright field pictures of an M-SCC control clone and a representative M-SCC *ZEB1* knockout clone (*ZEB1* KO #1), untreated (-TAM) or treated with Tamoxifen for 14 days (+TAM 14d). Scale bar: 100 µm. **e**, Bright field pictures of an E-SCC control clone and a representative E-SCC *ZEB1* overexpression clone (*ZEB1* OE #1), treated with DMSO or with 10 ng/ml, 100 ng/ml, or 1 µg/ml doxycycline (dox) for 7 days (+7d) or treated for 7 days followed by 14 days of withdrawal (+7d/-14d). Scale bar: 100 µm. **f**, Gating strategy for HMLE-Twist1-ER *ZEB1* overexpression and control cells.

